# Conformational analysis of membrane-proximal segments of GDAP1 in a lipidic environment using synchrotron radiation suggests a mode of assembly at the mitochondrial outer membrane

**DOI:** 10.1101/2023.05.08.539811

**Authors:** Aleksi Sutinen, Nykola C. Jones, Søren Vrønning Hoffmann, Salla Ruskamo, Petri Kursula

**Affiliations:** Faculty of Biochemistry and Molecular Medicine & Biocenter Oulu, University of Oulu, Oulu, Finland; ISA, Department of Physics and Astronomy, Aarhus University, Aarhus, Denmark; Department of Biomedicine, University of Bergen, Bergen, Norway

**Keywords:** Charcot-Marie-Tooth disease, GDAP1, mitochondrial outer membrane, protein biophysics

## Abstract

The mitochondrial outer membrane creates a diffusion barrier between the cytosol and mitochondrial intermembrane space, allowing the exchange of metabolic products, important for efficient mitochondrial function in neurons. The ganglioside-induced differentiation-associated protein 1 (GDAP1) is a mitochondrial outer membrane protein with a critical role in mitochondrial dynamics and metabolic balance in neurons. Missense mutations in the *GDAP1* gene are linked to the most common human peripheral neuropathy, Charcot-Marie-Tooth disease (CMT). GDAP1 is a distant member of the glutathione-S-transferase (GST) superfamily, with unknown enzymatic properties or functions at the molecular level. The structure of the cytosol-facing GST-like domain has been described, but there is no consensus on how the protein interacts with the mitochondrial outer membrane. Here, we describe a model for GDAP1 assembly on the membrane using peptides vicinal to the GDAP1 transmembrane domain. We used oriented circular dichroism spectroscopy (OCD) with synchrotron radiation to study the secondary structure and orientation of GDAP1 segments at the outer and inner surfaces of the outer mitochondrial membrane. These experiments were complemented by small-angle X-ray scattering, providing the first experimental structural models for full-length human GDAP1. The results indicate that GDAP1 is bound into the membrane *via* a single transmembrane helix, flanked by two peripheral helices interacting with the outer and inner leaflet of the mitochondrial outer membrane in different orientations. Impairment of such interactions could be a mechanism for CMT in the case of missense mutations affecting these segments instead of the GST-like domain.

## INTRODUCTION

Mitochondria can be distinguished from other cellular organelles by their dual membrane and the biochemical capabilities arising from endosymbiosis with ancient prokaryote features commonly referred to as holdover characteristics. They are solely responsible for eukaryotic cellular respiration and crucial for energy metabolism. They participate in exchanging metabolic products, regulation of reactive oxygen species (ROS) levels, and cellular calcium buffering.

In neurons, the importance of mitochondria arises from the immense energy consumption [1, 2]. Homeostasis both in the central nervous system (CNS) and peripheral nervous system (PNS) is closely regulated by the rate of oxidative respiration, enabling aerobic energy metabolism [3]. Moreover, the development and function of neurons, glial cells, and especially progenitor and neuronal stem cells are heavily impacted by mitochondrial function.

The mitochondrial membrane surrounds the organelle with the outer mitochondrial membrane (OMM), intermembrane space, and inner mitochondrial membrane (IMM), encapsulating the mitochondrial matrix. The membranes comprise similar composition in terms of lipid constituents with other cellular membranes. Mitochondrial membranes contain phosphatidylglycerol and cardiolipin (4–6). The phospholipid pool consists of phosphatidylcholine, phosphatidylserine, phosphatidylethanolamine, phosphatidylinositol and phosphatic acid. These lipids maintain the membrane potential and enable the membrane architecture required for respiration [4]. The tasks of the outer and inner membranes differ, as does their protein composition. The inner membrane is protein-rich and much more restrictive in terms of permeability, being saturated with proteins of the oxidative phosphorylation chain. The traffic between the mitochondrial matrix and the intermembrane space is driven by active transport. The outer membrane is much more permeable, allowing diffusion of metabolic products to and from the cytosol. Translocation of small globular proteins is possible through porous channelling protein complexes. The outer membrane enables contacts with other cellular compartments, such as the cytoskeleton, lysosomes, peroxisomes and endoplasmic reticulum. The interplay between other cell compartments and changes in mitochondrial morphology comprises mitochondrial dynamics, which couples mitochondrial fission and fusion. Specific proteins on the mitochondrial membranes, including the ganglioside-induced differentiation-associated protein 1 (GDAP1), participate in these events. Fission is driven by the soluble cytosolic dynamin-related protein 1 (DRP1), interacting with OMM partners, such as mitochondrial fission protein 1. Mitochondrial fission factors polymerize, creating a scission and separating the mitochondria [5]. Mitochondrial fission and fusion enable mitochondria to replicate and renew independently from the cell cycle. The detailed mechanism of GDAP1 in mitochondrial fission is unknown, but its role in the process has been demonstrated [6]. Consequently, mitochondrial dynamics directly affect neuronal function, and mitochondrial dysfunction is a leading cause of various neurodegenerative diseases, such as amyotrophic lateral sclerosis (ALS) and Charcot-Marie-Tooth syndrome (CMT).

Mutations in the *GDAP1* gene, expressed in neurons and glia to produce the OMM protein GDAP1, cause various subtypes of CMT, especially CMT4A and CMT2 [7–10]. The disease phenotypes feature fragmented and elongated mitochondria, leading to impaired fission and fusion [11–13]. In addition to diminished fission activity, lack of GDAP1 function leads to the accumulation of ROS, disturbance of Ca^2+^ buffering, and loss of lysosomal and cytoskeletal contacts [14–19]. Despite the GDAP1 effect on fission, its defined role, let alone the structural properties of the fission-fusion machinery, is unknown. GDAP1 is a member of the glutathione-S-transferase (GST) superfamily, but its possible enzymatic properties are unknown. Interestingly, gene homology within the GST family suggests that GDAP1 shares more similarities with prokaryotic than eukaryotic GSTs, which highlights its outlier role in the eukaryotic GST family and early divergence in evolution [20]. Previous structural studies [20–23] have focused on the two GST-like subdomains facing the cytosol, while only predictions exist for the structure of full-length GDAP1 on the OMM.

According to current understanding, GDAP1 is considered a classical tail-anchoring membrane protein (TA) with a single transmembrane helix. After the GST-like core domain, GDAP1 has a 60-residue C-terminus interacting with the OMM, which has been divided into two domains based on physicochemical properties: the hydrophobic domain (HD) and the transmembrane domain (TMD). The detailed topology or atomistic structure of the transmembrane region, or the membrane-vicinal segments, is not known.

We used synchrotron radiation circular dichroism spectroscopy (SRCD) to compare the secondary structures of GDAP1 membrane-proximal segments from both sides of the OMM in aqueous solution, membrane-like conditions, and lipid bilayer membranes. We then used oriented circular dichroism (OCD), where the peptide is embedded into macroscopically oriented bilayers to determine the average orientation of the peptide in the membrane. The method is based on Moffit’s theory [24], explaining the three transition dipole moments of the amide bond along either the perpendicular or the parallel axis. The transition of the dipole moment, *i.e.,* the axis with respect to the electromagnetic field vector, is monitored by inspecting the band shape at 208 nm [25], which allows to obtain information on peptide orientation with respect to the lipid bilayer plane. To complete the picture of full-length GDAP1 on the OMM, we additionally used small-angle X-ray scattering (SAXS) to study full-length human GDAP1.

## MATERIALS AND METHODS

### Synthetic peptides

The peptides for this study were purchased from Genscript Biotech (Rijswijk, Netherlands). The sequences of the peptides were selected using the structure prediction server TMHMM-2.0 [26] and the GDAP1 AlphaFold2 model [27]. The outer mitochondrial outer membrane peptide (OMOM) represents the outer leaflet part, ranging from residues 297-316 [NNILISAVLPTAFRVAKKRA]. The inner mitochondrial outer membrane peptide (IMOM) represents the inner leaflet tail-anchoring end, ranging from residues 341-358 [RKRLGSMILAFRPRPNYF]. Peptide concentration was adjusted to 1 mg/ml in deionized ultrapure water. The concentration was estimated based on the manufacturer’s data and verified on the AU-SRCD beamline based on UV absorbance.

### Liposome preparation

1,2-dimyristoyl-*sn*-glycero-3-phosphocholine (DMPC) and 1,2-dimyristoyl-*sn*-glycero-3-phospho-(1’-rac-glycerol) (DMPG) were purchased from Avanti Polar Lipids (Birmingham, AL, USA). Lipid stocks were prepared by dissolving dry lipid powder into methanol (DMPC) or an 80:20 chloroform-methanol mixture (DMPG). The concentration of each lipid stock was 10 mg/ml. Once completely dissolved, the lipids were mixed in 1:1, 2:1, 3:1 and 9:1 DMPC/DMPG molar ratios. The solvent was removed by evaporation in the hood under a gentle stream of pressurized air for 16 h. Small unilamellar vesicles (SUVs) were prepared by dissolving the dried lipid pellet into pure water at a final concentration of 15 mM. For freeze-thaw cycles, the lipid suspension was warmed in a +37 °C water bath, followed by snap freezing with liquid N_2_, for a total of seven times. The resulting multilamellar vesicles were then disrupted using sonication (Branson Model 450), 1 s on 2 s off, for 1 min at RT, followed by vigorous mixing until the vessel was completely clear. The resulting SUVs were stored at RT and used immediately for the experiment.

Bicelles were prepared by the addition of ultrapure water to the dried lipid pellets, followed by gentle agitation for 2 h at RT. The suspension was then treated with seven cycles of +37 °C (water bath) and -20 °C (slow freezing). Bicelle formation was induced by adding dodecyl phosphocholine (DPC) to a q-ratio of 2.85 [28]. The bicelles were stored at RT and used immediately for the experiment.

### Lipid film preparation for oriented samples

Sample preparation for the OCD experiment consisted of creating oriented lipid bilayers embedding the peptide molecules. The outline of the method used with peptides has been described previously [25, 29]. For all lipid-peptide films, we used 10 µg of peptide and 200 µg of lipid in ultrapure water.

The composition of the films is given in **Table 1**. The sample was distributed evenly onto a quartz plate (Suprasil, Hellma Analytics, Müllheim, Germany) and dehydrated at RT under a heating lamp (**Fig. 1B)**. The dehydrated films were assembled into a saturation chamber, containing a plate holder and a vessel containing a solution of saturated K_2_SO_4_ (**Fig. 1B)**. The relative humidity was monitored inside the chamber, which was between 85-89 % at +38 °C. The chamber was sealed, and the films were incubated for a minimum of 16 h, completing the sample preparation procedure (**Fig. 1A)**. The studied peptides represent the segments vicinal to the TMD (**Fig. 1C**).

**Table 1.** List of OCD lipid-peptide films and CD samples.

| **OCD** |  |
| --- | --- |
| **Film 1 (F1)** | 1:1 SUV DMPC/DMPG L/P 200:10 (w/w) |
| **Film 2 (F2)** | 2:1 SUV DMPC/DMPG L/P 200:10 (w/w) |
| **Film 3 (F3)** | 3:1 SUV DMPC/DMPG L/P 200:10 (w/w) |
| **Film 4 (F4)** | 1:1 bicelle DMPC/DMPG L/P 200:10 (w/w) |
| **Film 5 (F5)** | 3:1 bicelle DMPC/DMPG L/P 200:10 (w/w) |
| **Film 6 (F6)** | 9:1 bicelle DMPC/DMPG L/P 200:10 (w/w) |
| **SRCD** |  |
| **SUVs** | 1:1 DMPC/DMPG L/P 100:1 (mol/mol) |
|  | 2:1 DMPC/DMPG L/P 100:1 (mol/mol) |
|  | 3:1 DMPC/DMPG L/P 100:1 (mol/mol) |
|  | 3:1 DMPC/DMPG L/P 50:1 (mol/mol) |
|  | 3:1 DMPC/DMPG L/P 10:1 (mol/mol) |
| **CD (home source)** |  |
| **Detergents & other** | 5 mM DDM |
|  | 5 mM DPC |
|  | water |

**Figure 1.**
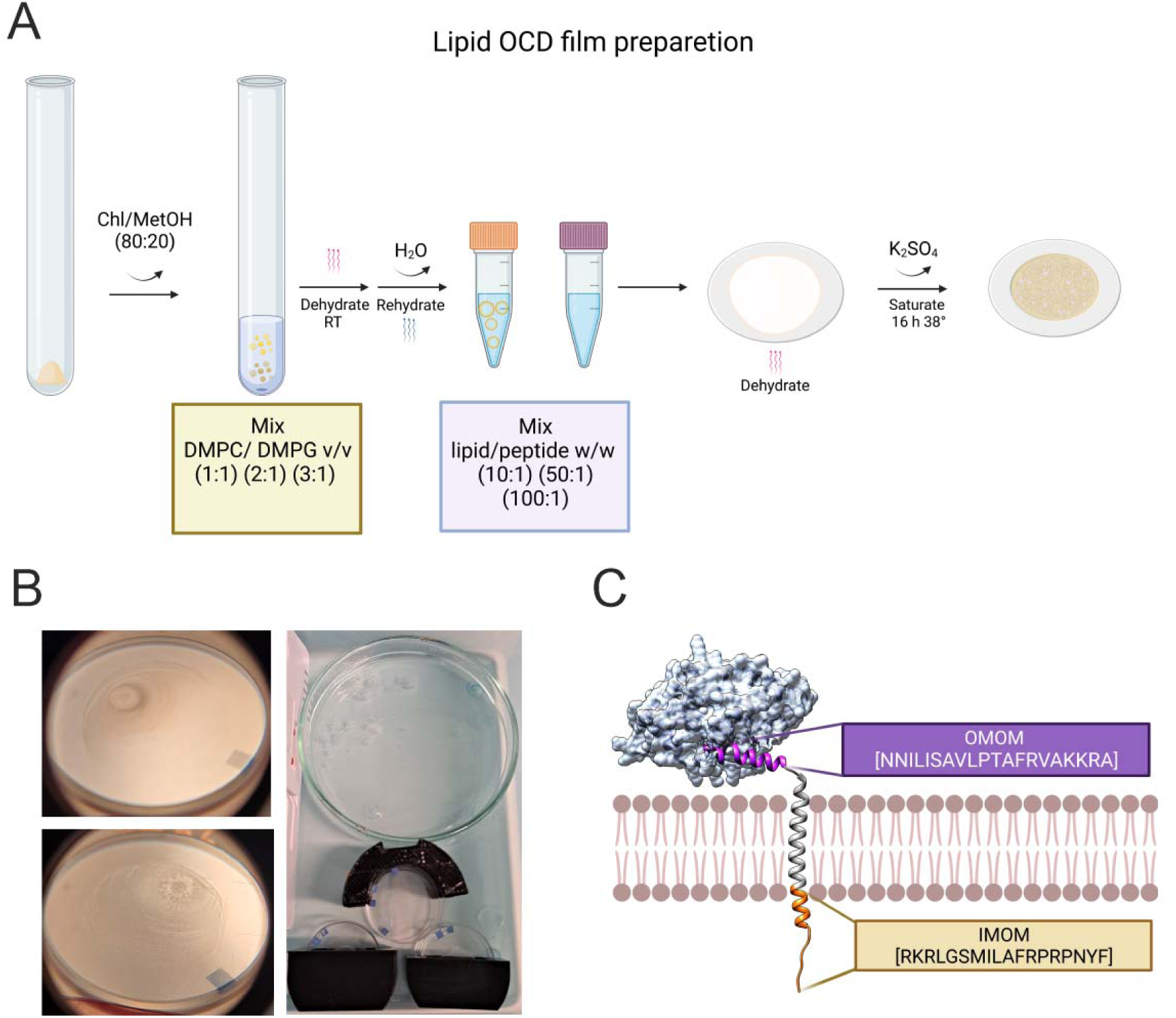
The experimental design of OCD. A. The liposomes are prepared in various composition ratios and the peptide are mixed in water. The mixture is dehydrated and then saturated with K_2_SO_4_ and incubated for> 16h before the measurement. B. The film should be homogenous and evenly distributed, and the saturation process must be done in a moderately isolated system with stable temperature and humidity. C. The regions of interest in GDAP1; the used peptides are highlighted. The figure was partially made using Biorender.

### Oriented and isotropic synchrotron radiation circular dichroism spectroscopy

Synchrotron radiation OCD and conventional SRCD spectra were collected on the AU-SRCD beamline at the ASTRID2 synchrotron (ISA, Aarhus, Denmark). Solution CD samples were prepared in ultrapure water, containing 0.2 mg/ml of peptide in 100:1, 50:1 and 10:1 molar lipid-to-peptide ratio (L/P). The SRCD samples were equilibrated to room temperature and applied into 0.2-mm closed circular quartz cuvettes (Suprasil, Hellma Analytics). SRCD spectra were recorded from 170 nm to 280 nm at +30 °C. Three scans per measurement were repeated and averaged. The complete set of CD and OCD samples is presented in **Table 1.**

The lipid-peptide films on the quartz plate were further assembled into a rotation sample holder. OCD scans were done at 90° intervals of rotation about the beam axis to complete a 360° rotation. Spectra were recorded from each position once and averaged. The spectra were processed using beamline software and CDToolX [30].

The samples containing detergent micelles contained 5 mM of *n*-dodecyl-β-D-maltopyranoside (DDM) or 5 mM DPC. The detergent CD spectra were collected using a Chirascan CD spectropolarimeter (Applied Photophysics, Surrey, UK), using a closed 1 mm quartz cuvette (Hellma Analytics). The data were collected between 185-280 nm.

### Small-angle scattering

Full-length GDAP1 was cloned into the pFastBac-Dual HT vector. An N-terminal His_6_-tag and a Tobacco Etch virus protease (TEV) digestion site were inserted ahead of GDAP1.

The construct was transformed into the *E. coli* DH10Bac strain for bacmid generation. Bacmids were purified, and 2 µg of bacmid DNA were used to transfect *Sf21* cells for baculovirus generation using FuGENE® transfection reagent (Promega). The protein was expressed in Lonza Insect-EXPRESS™ medium for 72-96 h, 100 rpm, +27 °C. The cells were washed in phosphate-buffered saline (8 mM Na_2_HPO_4_, 2 mM KH_2_PO_4_, 137 mM NaCl, 2.7 mM KCl (pH 7.4) (PBS)) and snap frozen with LN_2_.

The frozen cells were re-suspended into 40 mM HEPES pH 6.88, 400 mM NaCl, 10% glycerol, 25 mM imidazole, 1 x EDTA-free protease inhibitor (Sigma). Lysis was done using a glass douncer on ice with 50-70 revolutions. DNA was disrupted by incubating the lysate on ice with 0.2 mg/ml DNase I (Applichem) and 0.2 M CaCl_2_ in lysis buffer. The supernatant was clarified at +4 °C, 1 h, 234788 g, and the supernatant was discarded. The membrane fraction pellet was resuspended into 1% (w/v) DDM (∼195 mM) or DPC (∼284 mM) in lysis buffer. The membrane fraction was solubilized at +4 °C, 2 h with continuous rotation. The membrane lysate was clarified by ultracentrifugation at +4 °C, 1 h, 234788 g. The solubilized membrane fraction was passed through HisPur® (Thermo Fisher)-affinity resin by gravity flow. The resin was washed with 50 mM imidazole, and elution was done with 300 mM imidazole pH 6.88 in lysis buffer. The fractions were collected and verified with SDS-PAGE and MALDI-TOF/MS (Bruker UltrafleXtreme). The samples were dialyzed using a 14 MWCO dialysis membrane in 25 mM HEPES pH 6.88 (RT), 300 mM NaCl, 10% (w/v) glycerol and 2 mM DPC. TEV protease was added with a 1:5 molar ratio to cleave fusion tags. Reverse IMAC was used to separate the cleaved protein by gravity flow.

Samples were concentrated and injected into Superdex 200 10/300 column (S200) (cytiva) using 25 mM HEPES pH 6.88 (RT), 300 mM NaCl and 2 mM DPC as mobile phase (SEC buffer). SEC-SAXS data were collected on the SWING beamline (SOLEIL, Paris) [31], using a 0.5 ml/min flow rate. The data acquisition from the SEC peaks was done with a rate of 3 s/frame. The SAXS data were reduced using Foxtrot-3.5.10 and ATSAS [32].

### Structure prediction and modelling

3D structures of the IMOM and OMOM peptides were predicted using the PEP-FOLD server [33]. The top-ranked model of each was then modelled onto a DMPC bilayer with PPM [34], using the default parameters of the server. The AlphaFold2 model of human GDAP1 was modelled into the OMM using CHARMM-GUI [35]. The composition of the membrane is available on the CHARMM-GUI website (https://charmm-gui.org/?doc=archive&lib=biomembrane). The disordered N terminus (23 residues) had to be removed from the model to align the TM helix properly with the membrane plane. Electron density maps of GDAP1 were computed using the full-length GDAP1 scattering profiles using DENSS. The parameters of the algorithm, map averaging and sharpening can be found in the original publication [36].

## RESULTS AND DISCUSSION

Having determined the crystal structure of the GST-like core domain of GDAP1 earlier (18, 19), to shed light on the missing parts of the full-length GDAP1 structure, we carried out SRCD experiments on two membrane-proximal peptides, located immediately before and after the GDAP1 TMD. Experiments were carried out both in an isotropic setting in solution and using oriented rehydrated membranes in OCD, to follow peptide folding under different membrane-mimicking conditions and observing peptide orientation with respect to the membrane surface.

### CD spectroscopy of isotropic samples in solution

Both the IMOM and OMOM peptides were soluble in water. Folding of both peptides was observed when DMPG/DMPC SUVs were mixed with the peptides (**Fig. 2)**. In these samples, we saw a high positive peak at 195 nm and a minimum at 208 nm, followed by another minimum at 222 nm. Hence, the SRCD spectra show that both peptides gain α-helical structure, when mixed with liposomes, but the spectral shapes were different. To investigate the required lipid-to-peptide ratio for the peptide to fold, a stepwise increase of lipid was done, from 10:1 to 50:1 and 100:1 L/P ratio (**Fig. 2B, D)**. The lowest ratio does not contain enough lipids per peptide molecule to induce folding. With a five-fold increase to 50:1, a positive 195 nm peak appears. The IMOM negative peaks clearly distinguish at 208 nm and 222 nm, but in OMOM, they are less well defined. When the ratio increases to 100:1 L/P, both IMOM and OMOM have strong α-helical spectra with well-defined peaks; however, their shapes are different, indicating non-identical structures for the two peptides when membrane-bound. The difference in the ratio of DMPG/DMPC, *i.e.* the surface charge of the membrane, had no effect on the shape of the spectra, when membranes with 25% or 50% negatively charged DMPG were compared (**Fig. 2A, C)**.

**Figure 2.**
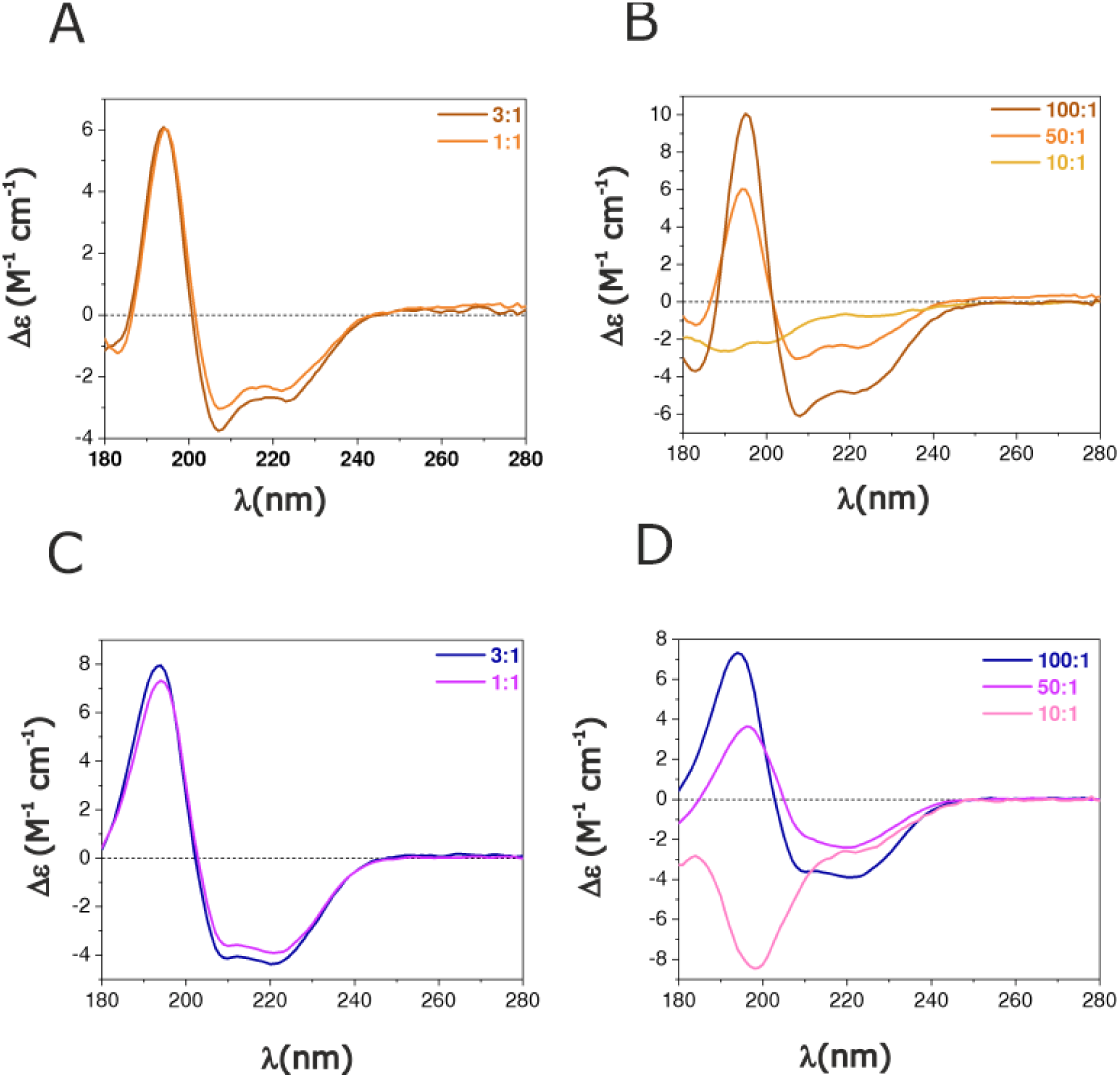
Isotropic SRCD measurements. Spectra for IMOM (A-B) and OMOM (C, D). The peptides folded in DMPC/DMPG-based SUVs. A and C show a comparison of different DMPC/DMPG ratios at a 100:1 L/P ratio; the DMPC/DMPG ratio did not have an effect. B and D compare different L/P ratios at a DMPC/DMPG ratio of 1:1. Both peptides required more than a 10:1 L/P ratio to fold.

### Oriented CD spectroscopy of rehydrated samples

OCD was measured from oriented, rehydrated lipid-peptide multilayer films to determine the average orientation of the helical peptide structure in the lipid bilayer. The films were prepared according to **Table 1 and Fig. 1.** The films were prepared in the same batch as the background film without the peptide to combat sources of heterogeneity, such as unevenly distributed samples, relative humidity, and temperature. The mixture spreading into the film was consistent throughout the experiment. To monitor possible light scattering and linear dichroism effects, OCD spectra were collected from 160 nm to 280 nm at different angles about the beam axis. The absorbance curves from each angle showed reproducibility, and there was slight variation between each scan in all films (**Fig. S1, S2)**, which is normal in OCD experiments. No interference from the film or signs of oversaturation were observed.

#### The IMOM peptide in an oriented environment

A qualitative comparison of the IMOM OCD spectra shows that the peptide lies on the plane of the membrane, when 50% of the headgroups have negative charge. The conformation then tilts slightly, when the negative charge on the membrane is decreased. Compared to the solution CD spectrum, which shows that the peptide is helical in a lipidic environment embedded in aqueous solvent, the positions of the CD peaks do not move (**Fig. 3)**. This indicates similar folding in isotropic and oriented samples.

**Figure 3.**
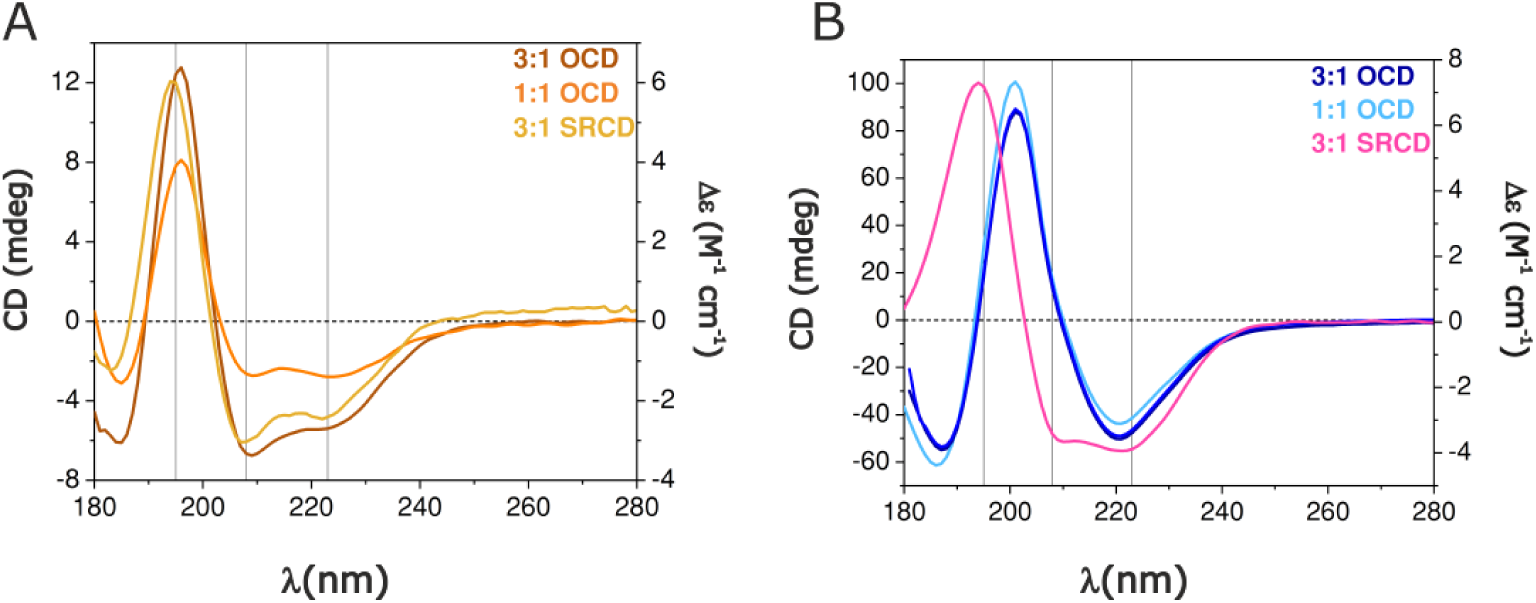
Comparison of OCD and isotropic SRCD. A. OCD spectra recorded for IMOM in orange traces compared with 3:1 DMPC/DMPG isotropic spectrum in yellow trace. B. For OMOM, the OCD traces are shown as navy and cyan and SRCD in magenta. The three vertical lines highlight the 195-, 208- and 222-nm peaks.

An important aspect of OCD is that analysis generally focuses on the spectral shape, rather than the exact amplitudes. Aggregation of the peptide-lipid complexes or uneven distribution of sample can alter the peak heights. The sample distribution was good, based on the absorbance spectra measured for each scan (**Fig. S1)**. Especially for l7-helical peptides, one wants to follow if the peak positions move, and if the ratio between the 208-nm and the 222-nm peak changes, which gives indications of helix orientation. The shape of the IMOM OCD spectra showed that the 208 nm CD peak was well defined, being strongest at the DMPC/DMPG ratio of 1:1. Its ratio to the 222-nm peak then decreases upon loss of negative membrane charge, indicating tilting of the helix (**Fig. 3A)**. Films made of bicelles gave essentially the same result as regular lipid bilayers, indicating that bicelle samples could be similarly oriented for OCD experiments as regular bilayers (**Fig. 4A)**.

**Figure 4.**
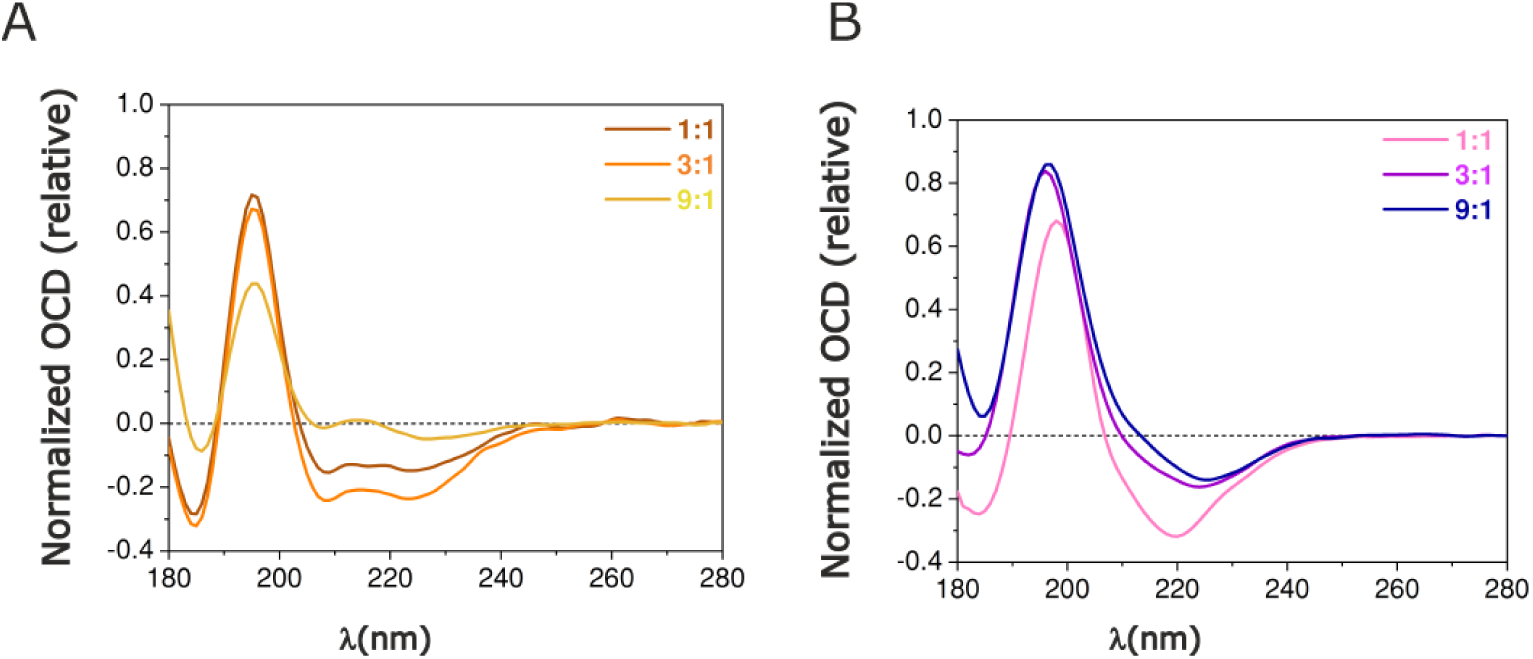
Normalized OCD spectra from bicelle films at different DMPC/DMPG ratios. A. IMOM. B. OMOM. The molar lipid-to-protein ratio was 100:1.

In summary, both in isotropic SRCD and OCD, the IMOM peptide has the spectral shape of a characteristic α-helix. The significant negative 208-nm CD peak with respect to the 195-nm and 222-nm peaks in the OCD experiment suggests that most of the peptide bonds lie parallel to the electric field, or perpendicular to the direction of propagation of light [25]. The shape of the spectra suggests that the IMOM peptide is lying flat on the lipid surface, and that it can be induced to tilt slightly by changing the lipid composition. Decreasing the negative charge of the membrane surface appears to induce tilting of the peptide, which appears parallel to the membrane surface in 1:1 DMPC/DMPG.

#### The OMOM peptide in an oriented environment

The magnitude of the CD signal from the films with OMOM was much higher than with the IMOM peptide, averaging ∼20 mdeg and exceeding 50 mdeg (**Fig. 3B),** which could indicate differences in sample coverage or peptide aggregation state on the membranes. The OCD curve overall shape largely lacks the negative minimum at 208 nm, which is an indication of a tilted or even transmembrane conformation [25, 37]. The isotropic SRCD spectra for OMOM have a weaker 208-nm band compared to IMOM, which indicates specific differences in the two peptides; the 208-nm band for OMOM is more like a shoulder to start with, before orienting the bilayers. In comparison to the isotropic spectra with SUVs or bicelles, the 208-nm band is nearly missing in OCD, while the 195-nm band moves towards a higher wavelength in some samples, to 198-204 nm (**Fig. 4B)**. This is a normal phenomenon for OCD of a tilted peptide [25, 37]; as the 222-nm peak does not move, the peptide does not transition towards β-stranded structure. There were no peak shifts when the peptides bound to SUVs were compared to the bicelle conditions.

The results do not support the hypothesis that the OMOM peptide is parallel to the membrane surface, as predicted by the AlphaFold2 model inserted into the OMM membrane composition (**Fig. 1C, S3)**. The overall shape of the spectra and weakness of the 208 nm peak suggest that some fraction of the peptides are buried in the lipid membrane, possibly in a highly tilted conformation. The samples may contain a mixture of orientations, and the results reflect an average of these. As opposed to IMOM, membrane surface charge does not affect the apparent orientation of OMOM. It should be noted that the OMOM peptide carries a Pro residue in the middle region, and PEP-FOLD predicts either a kinked helix or two short helices with an angle between them. This could enable tilting of the peptide in a similar way as earlier observed for a membrane-binding peptide from the myelin protein P0 cytoplasmic domain [37], which showed a similar OCD spectrum.

### Small-angle X-ray scattering of full-length GDAP1

The results from the SRCD and OCD experiments showed that the GDAP1 segments close to the TM helix bind to lipids. We then investigated whether the full-length recombinant GDAP1 could be isolated from membranes and purified. To select the detergent, CD spectra of IMOM and OMOM were measured to compare their folding in water, DDM and DPC. The CD spectra showed that both peptides have no secondary structure features in water. When DDM was used, the spectra did not show a clear difference compared to the spectra measured in water. A dramatic difference was seen when DPC micelles were added; folding of both peptides into l7-helical structures was observed, similarly to the experiments with liposomes above (**Fig. 5A)**. The effect of DPC is logical due to the similarity of its headgroup to DMPC (**Fig. 5B)**. The importance of the lipid headgroup is highlighted by the fact that the peptides did not fold in DDM but became helical in DPC. DPC was therefore deemed suitable to stabilize the protein *in vitro* and used for the purification (**Fig. 5C)**.

**Figure 5.**
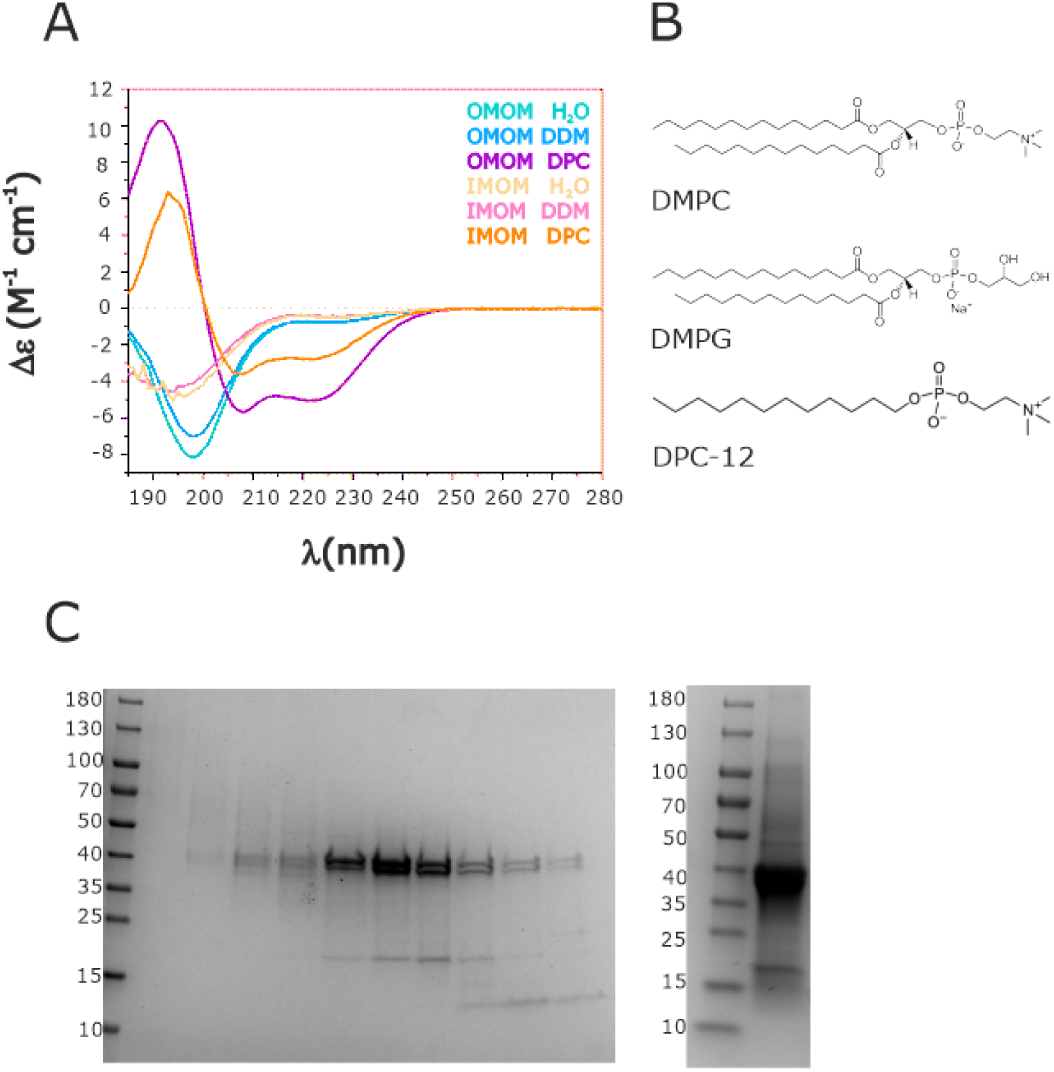
Detergent selection for full-length GDAP1 purification. A. Isotropic CD measurements for the IMOM and OMOM peptides in water and detergent micelles. B. Molecular structures of DMPC, DMPG, and DPC. Note the similarity between the headgroups of DMPC and DPC. C. Full-length GDAP1 purification after SEC. Left: Fractions collected from the main eluting peak in SEC. Right: concentrated sample pooled from the main peak and further used for SEC-SAXS.

SEC-SAXS data were collected from the purified full-length GDAP1 samples in DPC buffer. The SEC-SAXS retention profile shows two distinct peaks, based on further analyses likely corresponding to GDAP1 with and without micelle (**Fig. 6, S4, S5, S6**). The SAXS data are summarized in **Table 2.**

**Figure 6.**
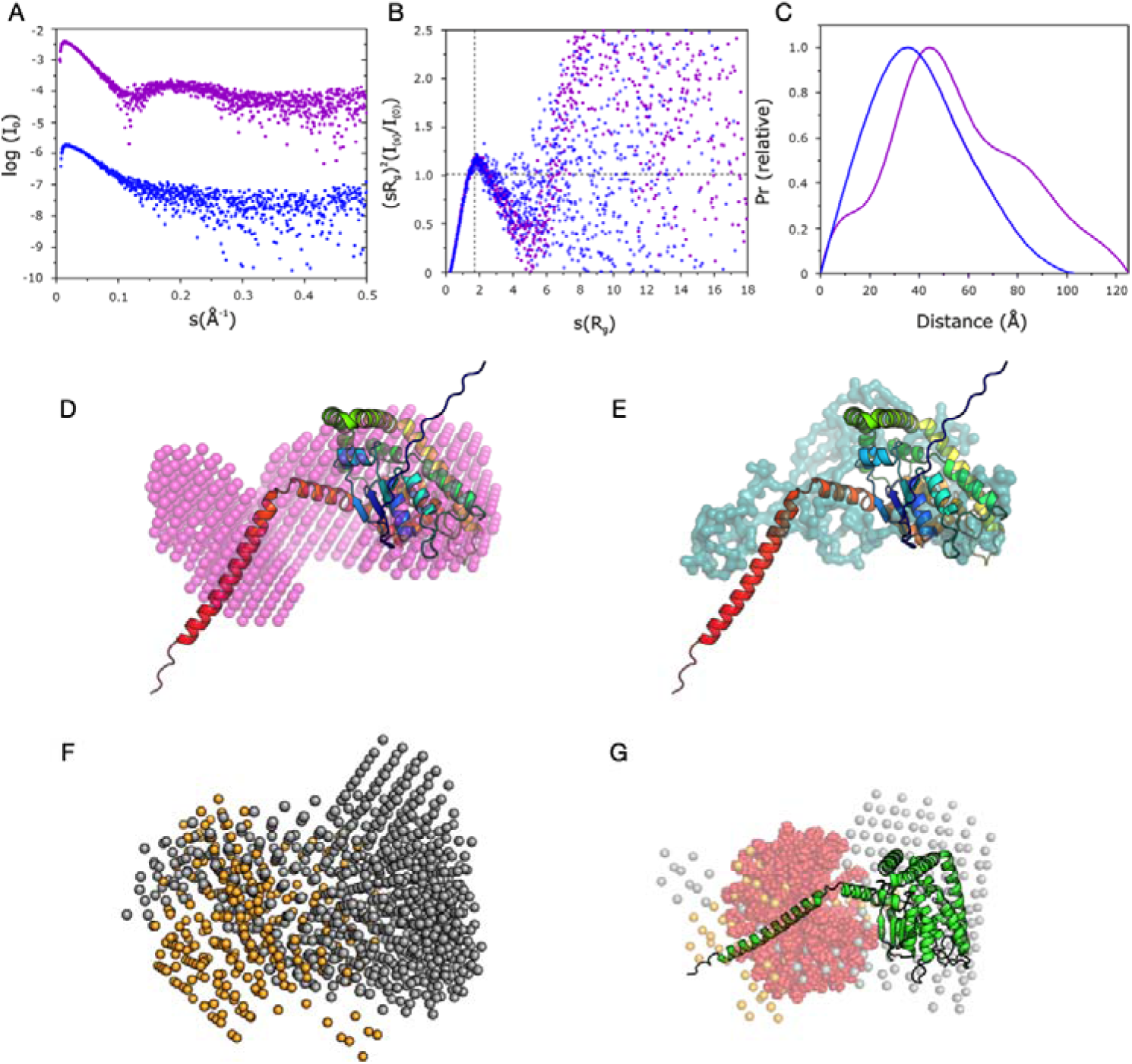
SAXS analysis of full-length human GDAP1. A. Scattering curves for the two SEC-SAXS peaks for full-length GDAP1 in DPC. The SAXS data are coloured purple (first peak) and blue (second peak). B. Dimensionless Kratky plots for the samples indicate similar levels of flexibility. C. Distance distribution indicates that the first peak has a particle with shoulders characteristic of a detergent micelle, while the second peak corresponds to a folded protein without bound micelle. D. DAMMIN model (magenta) for the monomeric full-length human GDAP1 superimposed with the AlphaFold2 model (cartoon) indicates presence of the GST-like domain to the right and the C-terminal region including the TMD to the left. E. Chain-like GASBOR model for the monomer. F. MONSA models for GDAP1 with a DPC micelle. 4 models are superposed, resulting from separate runs with different contrast values for the micelle. The protein phase is shown as grey beads and the micelle phase is in orange. G. Superposition of one MONSA model with a model of GDAP1 (cartoon) bound to a DPC micelle (red). Fits of the various models to the experimental data are in Fig. S6.

**Table 2.** SEC-SAXS experiments on full-length GDAP1 in DPC-containing buffer. The CRYSOL fits correspond to the AlphaFold2 model.

| Peak | 1<sup>st</sup> peak (containing micelle) | 2<sup>nd</sup> peak (detergent-free) |
| --- | --- | --- |
| R<sub>g</sub> (Guinier) (Å) | 42.06± 1.38 | 31.53± 0.54 |
| R<sub>g</sub> (GNOM) (Å) | 42.06 | 31.56 |
| D<sub>max</sub> (Å) | 113.36 | 102.71 |
| V<sub>p</sub> (Å<sup>3</sup>) | 220212 | 101981 |
| Estimated MW Bayes/V<sub>c</sub> (kDa) | 94.2/52.2 | 41.9/42.7 |
| χ<sup>2</sup> (GASBOR) | - | 0.95 |
| χ<sup>2</sup> (DAMMIN) | - | 0.95 |
| χ<sup>2</sup> (CRY SOL) | 6.41 | 2.11 |

A comparison of the SAXS scattering profiles shows that the first eluting peak contains a protein and a micelle together. However, the peak eluting later does not seem to contain a micelle, suggesting that full-length GDAP1 can remain soluble also in the absence of detergent in micellar form. This was earlier seen for the closest homologue of GDAP1, GDAP1L1 [22], which also has the HD and TMD domains. Moreover, the R_g_ and estimated MW of the peaks suggest that the scattering arises from a monomeric GDAP1. The R_g_ and D_max_ of the GDAP1 AlphaFold2 model are 31.1 and 93 Å, respectively; the values are close to those for the second eluted peak, further indicating the presence of a monomer. *Ab initio* bead-based modelling of the structure results in an elongated envelope that fits the GST-like domain, with an additional density corresponding to the size and location of the HD and TMD (**Fig. 6**). The same observation was concurred by computing the 3D electron density map from the solution scattering profile using DENSS (**Fig. S4, S5**). The volume corresponds to a monomer, which may indicate that either the full-length GDAP1 does not form dimers, as shown for constructs lacking the C terminus [20, 22, 23], or the concentration in the experiment was so low that monomers were predominant. The monomer-dimer equilibrium of GDAP1 *in vitro* is dependent on the protein concentration [20]. There is no evidence for the presence of a detergent micelle in the particle eluting in the second peak, as also indicated by the distance distribution profile. The results show that full-length GDAP1 is soluble without a bound detergent micelle. Detergent monomers may be bound to the hydrophobic surfaces in the experiment, but not to the extent that would cause characteristic minima in the SAXS profile.

### Implications for full-length GDAP1

Membrane interactions are important for protein function and localization. Typically, the residues interacting with lipid membranes fold into helices. The helix confirmation may vary based on the amino acid sequence and lipid composition. OCD is a technique that extends the conformational information obtained from CD spectroscopy to address the average orientation of the polypeptide in the membrane structure.

The crystal structure of the human GDAP1 cytosolic GST-like domains, ranging to residue 302, contains a unique dimer interface [20, 22, 23]. GDAP1 has two domains following residue 302, the HD and TMD [8, 9]. OMM proteins, many being fission factors, have a TA membrane-binding domain [38]. The TA translates the protein from the endoplasmic reticulum (ER) into mitochondria and docks it into the outer membrane [39]. Immunohistochemical tracking has demonstrated that HD and TMD domains are sufficient for mitochondrial localization and fission activity of GDAP1 [40]. The critical differences between CMT-linked missense point mutations in the HD and TMD [41–43] suggest that the residues before residue 320 have more to do with the fission function, and the remaining C-terminal tail is for targeting GDAP1 to the OMM.

The experimental scheme in this study presents a model of the GDAP1 TA segment, and our SAXS data provide the first experimental models for full-length human GDAP1. The peptide OMOM represents the HD domain, and IMOM represents the TMD tail in the intermembrane space. There was no prior evidence of either being inserted into the membrane as a transmembrane helix. SRCD spectra of both peptides showed helical conformation when bound to lipid bilayers, and OCD data displayed different conformations. The IMOM peptide was more parallel to the membrane surface, while OMOM was more tilted into the membrane. Modelling of the AlphaFold2 structure of GDAP1 onto the OMM (**Fig. S3**) supports these findings, showing that both peptides must be interacting with the membrane surface. A distinguishing factor is that the outer leaflet helix is likely more flexible with many conformations, including being partially buried. In contrast, the inner leaflet helix acts as a fixed anchor following the transmembrane helix.

The OMOM peptide is a site for CMT disease mutations; known mutations include R310Q and R310W, which we characterized before [20], as well as N297T and P306L [44, 45]. Arg310 is predicted by AlphaFold2 to interact with the GST-like core domain, pointing away from the membrane [20]. Therefore, the OMOM segment or the HD is likely a structure that links the membrane surface to the folded core domain on the cytosolic side, and mutations interfering with this linkage may cause disease. Co-sedimentation experiments suggest that the helix contributes to membrane binding (unpublished data). The IMOM peptide, *i.e.* the GDAP1 C-terminal tail in the mitochondrial intermembrane space, is a target for at least two known CMT mutations: L344R and S346N [46, 47]. Leu344 is likely embedded in the membrane, and mutation to Arg would interfere with this scenario, possibly affecting GDAP1 targeting. Predicted models for the two peptides are shown in **Fig. 7**, further supporting their different folding properties and membrane interactions.

**Figure 7.**
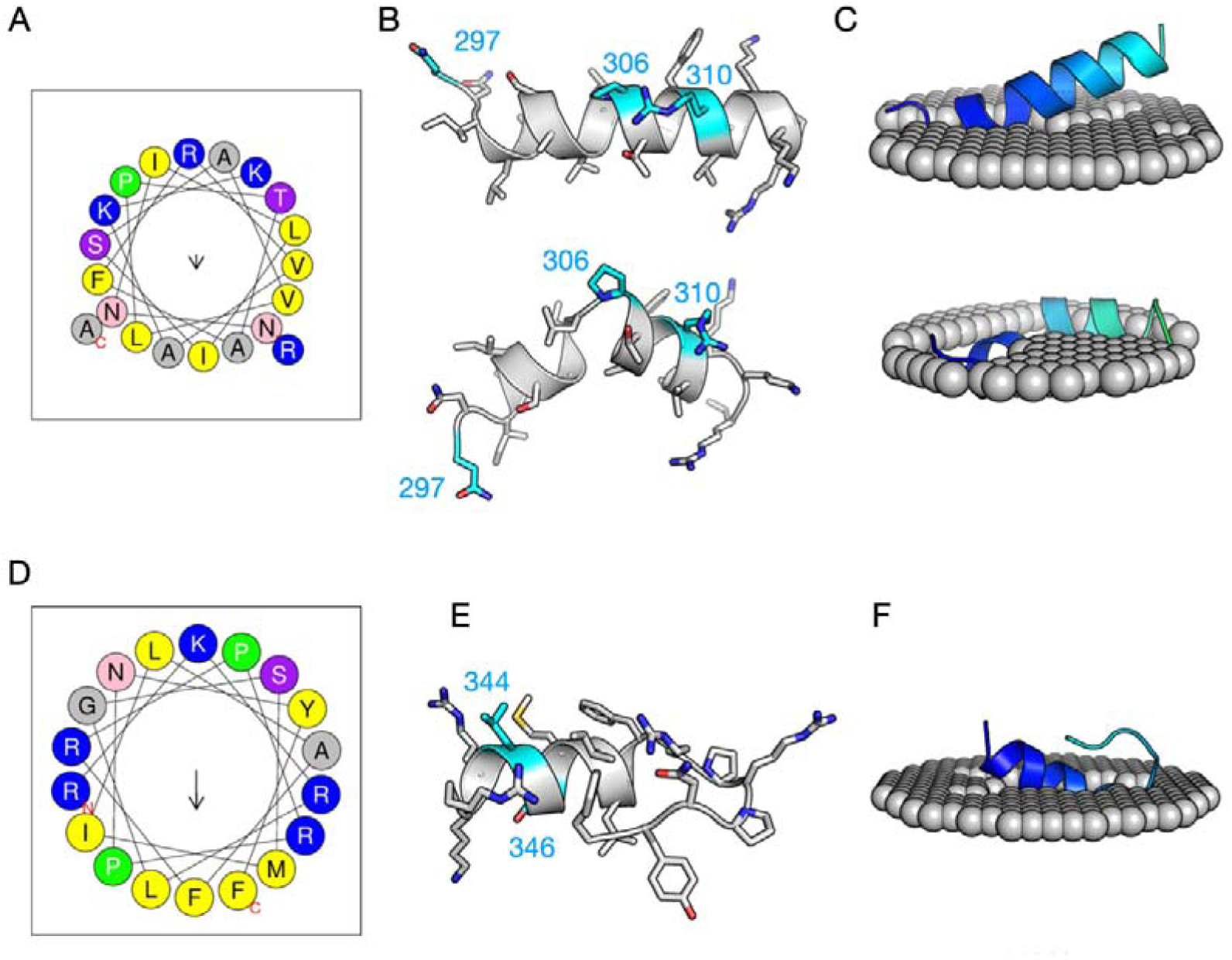
Peptide structure predictions. A. Helical wheel diagram of OMOM. B. Two predicted conformations of OMOM from PEPFOLD. The locations of CMT mutation sites (Asn297, Pro306, Arg310) are indicated in cyan. C. PPM docking of the two OMOM models onto a DMPC membrane. D. Helical wheel diagram of IMOM. E. Predicted conformation of IMOM. CMT mutation sites (Leu344, Ser346) are indicated in cyan. F. PPM docking of the IMOM model onto a DMPC membrane.

Based on current experimental data and computational predictions, the topology of the GDAP1 membrane interaction is a single transmembrane helix, with two flanking peripheral helices in both the OMM inner and outer leaflets. It should be noted that the system investigated here consisted of simplified liposomes and peptides, both mimetics of the interaction between GDAP1 and the OMM. The complete composition of the mitochondrial outer membrane is much more complex [48], and high-resolution structural studies on full-length GDAP1 in a physiological membrane lipid composition remain to be performed.

## Conclusions

Our results describe the OMM binding contributions of the flanking helices adjacent to the GDAP1 transmembrane helix. The IMOM and OMOM helices are not transmembrane but aligned along the membrane surface or tilted and partially embedded. The C-terminal tail in the intermembrane space is, thus, not disordered but tightly coiled into the inner leaflet, locking the protein into the OMM. The residues adjacent to the outer leaflet may have less affinity to the membrane, allowing more flexibility. These membrane-proximal helices of GDAP1 contribute to OMM binding and, thus, likely have a role in the localization, conformation, and function of GDAP1. The results further shed light on the potential disease mechanisms in CMT, whereby the membrane interactions of the segments outside the GST-like domain may be impaired, leading to abnormal protein targeting and membrane protein complex assembly.

## ACKNOWLEDGEMENTS

This work was funded by the Academy of Finland, project number 24302881. This project has received funding from the European Union Horizon 2020 research and innovation programme under grant agreement 101004806; beamtime on the AU-SRCD beamline was allocated *via* the MOSBRI transnational access proposal ID MOSBRI-2022-76. We wish to thank Dr. Arne Raasakka for the pilot development of OCD experiments on the AU-SRCD beamline prior to the current work. Beamtime and support on the SOLEIL synchrotron are gratefully acknowledged.

---

**Figure S1.**
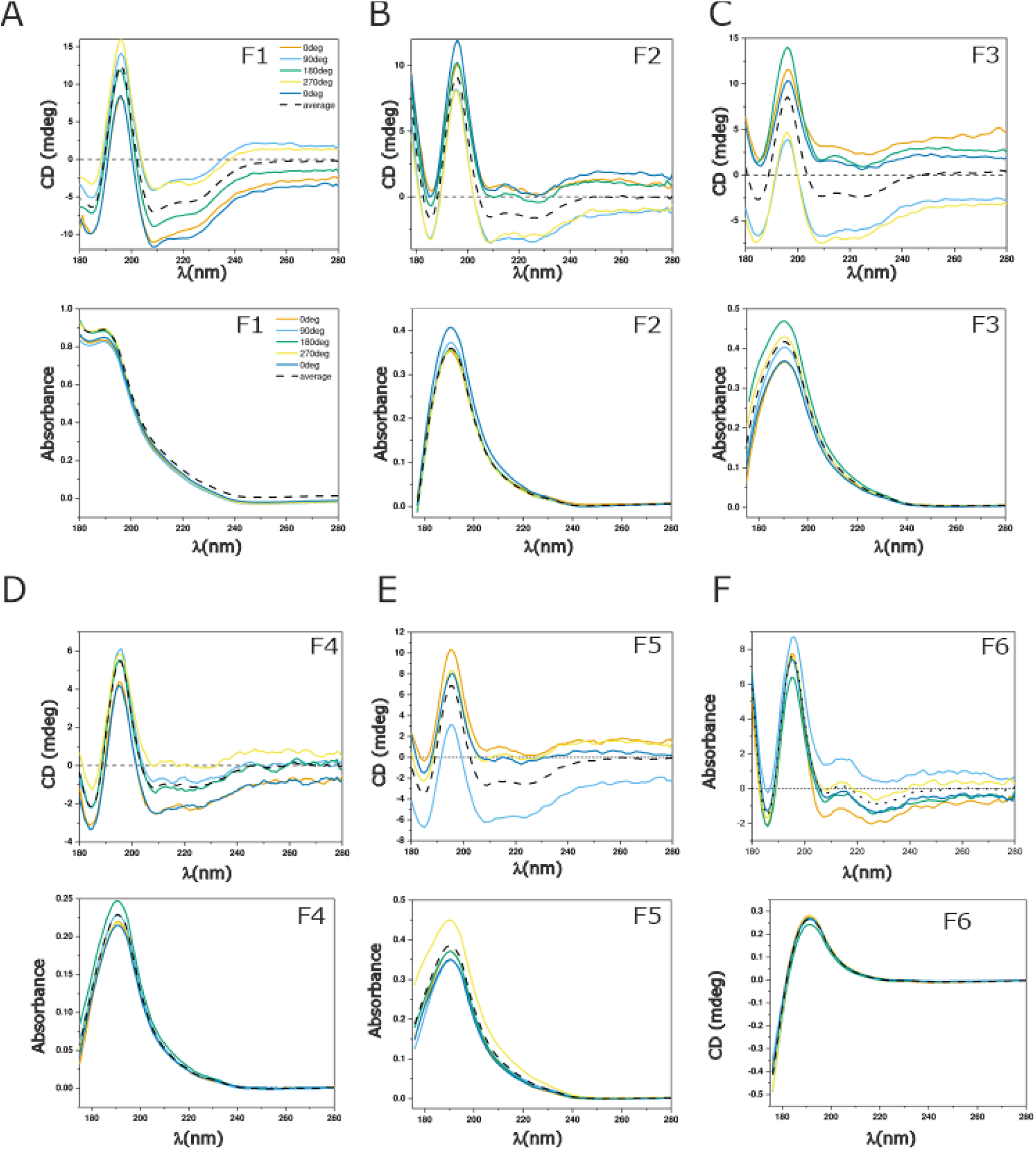
Complete OCD measurements from DMPC/DMPG films with the IMOM peptide. The averaged spectra are comprised of a full 360° turn in 90° intervals at +30 °C. A-F, data for film compositions F1-F6, respectively. CD signal of each interval (top) and corresponding absorbances (bottom) are shown. See Table 1 for film composition.

**Figure S2.**
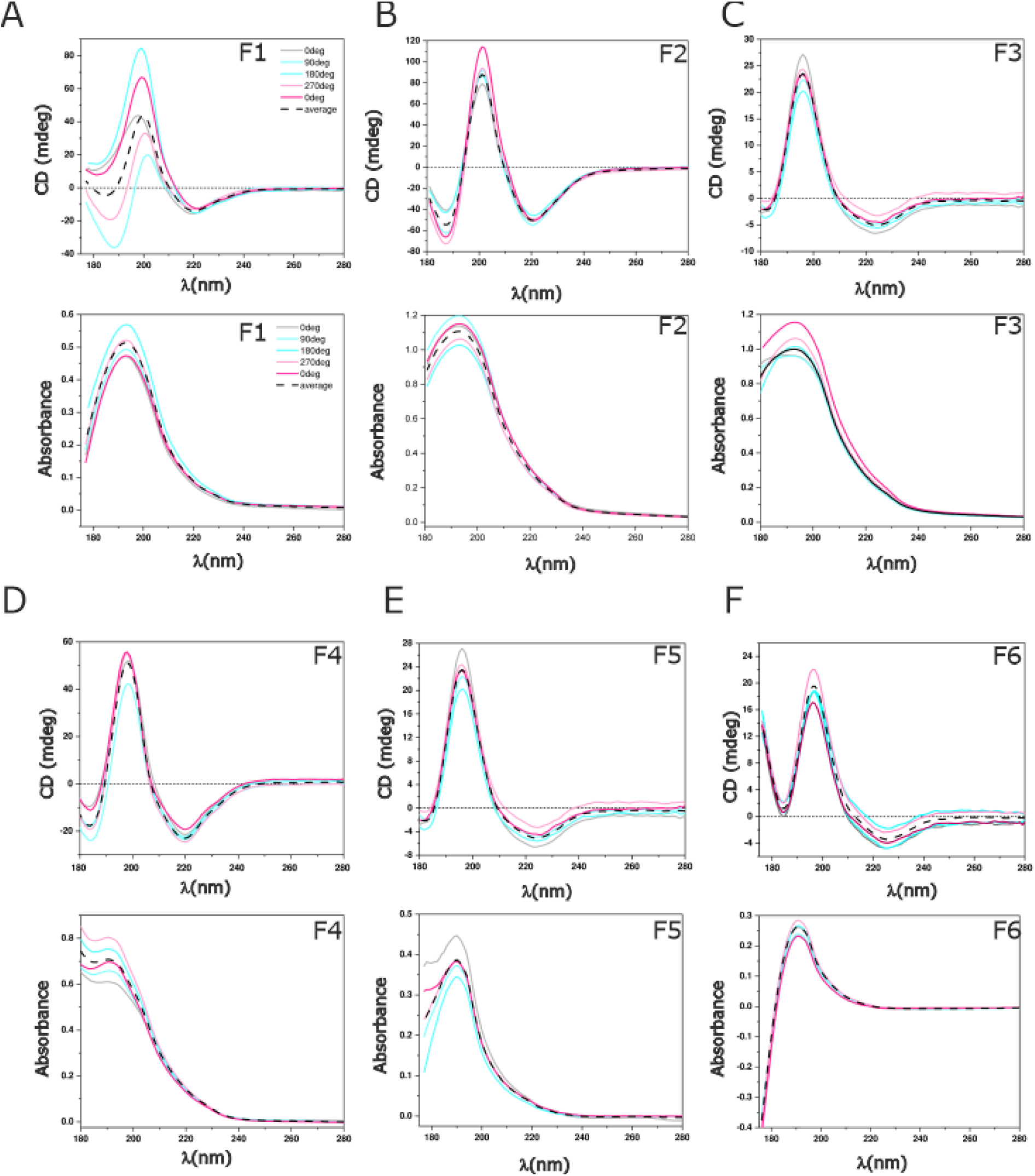
Complete OCD measurements from DMPC/DMPG films with the OMOM peptide. The averaged spectra are comprised of a full 360° turn in 90° intervals at +30 °C. A-F, data for film compositions F1-F6, respectively. CD signal of each interval (top) and corresponding absorbances (bottom) are shown. See Table 1 for film composition.

**Figure S3.**
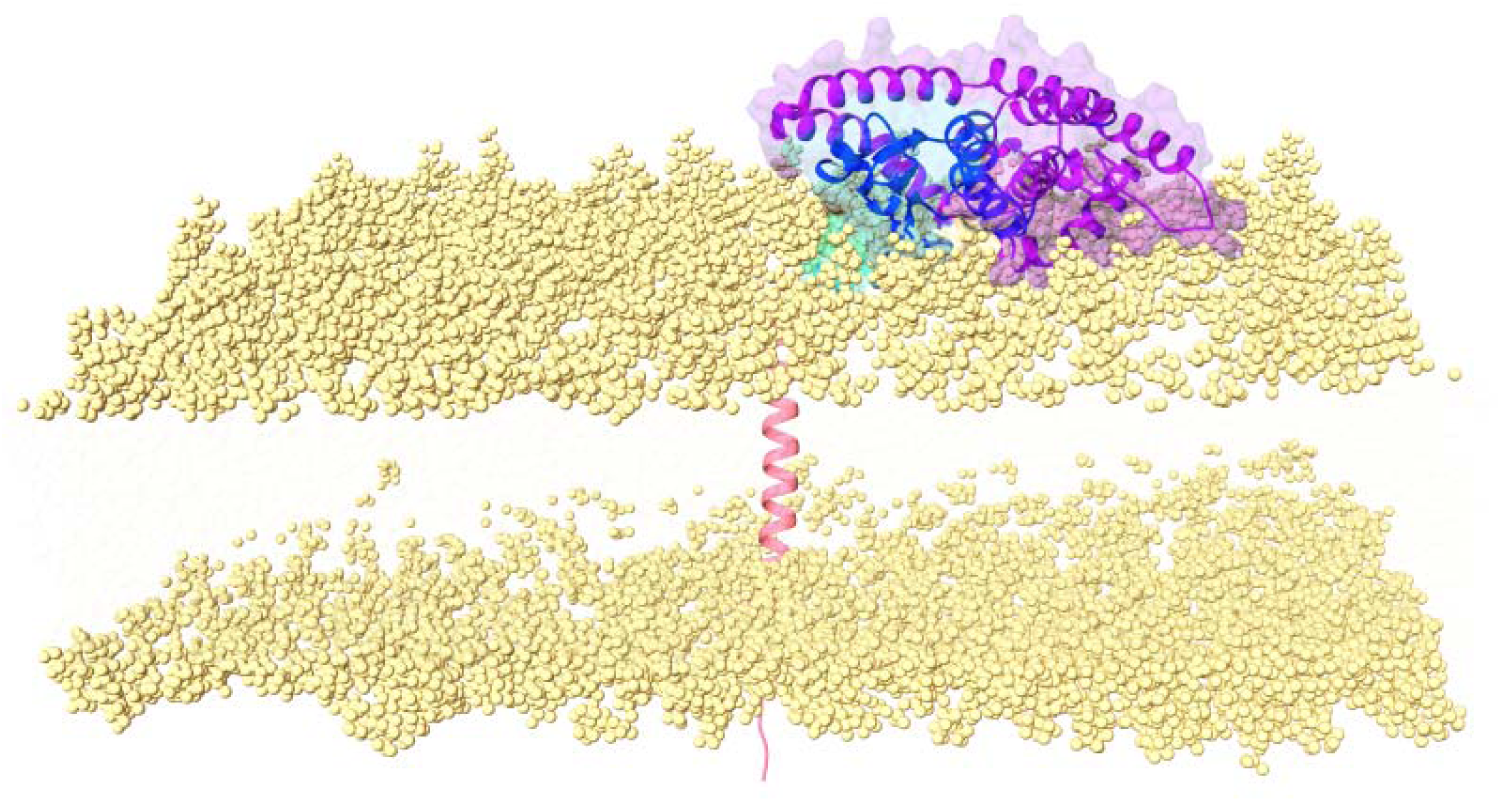
GDAP1 AlphaFold2 model inserted into the OMM membrane using CHARMM-GUI. The OMOM peptide is between the folded GST-like domain and the TMD, while the IMOM peptide is C-terminal to the TMD on the other side of the membrane. Lipid headgroups are shown as yellow spheres.

**Figure S4.**
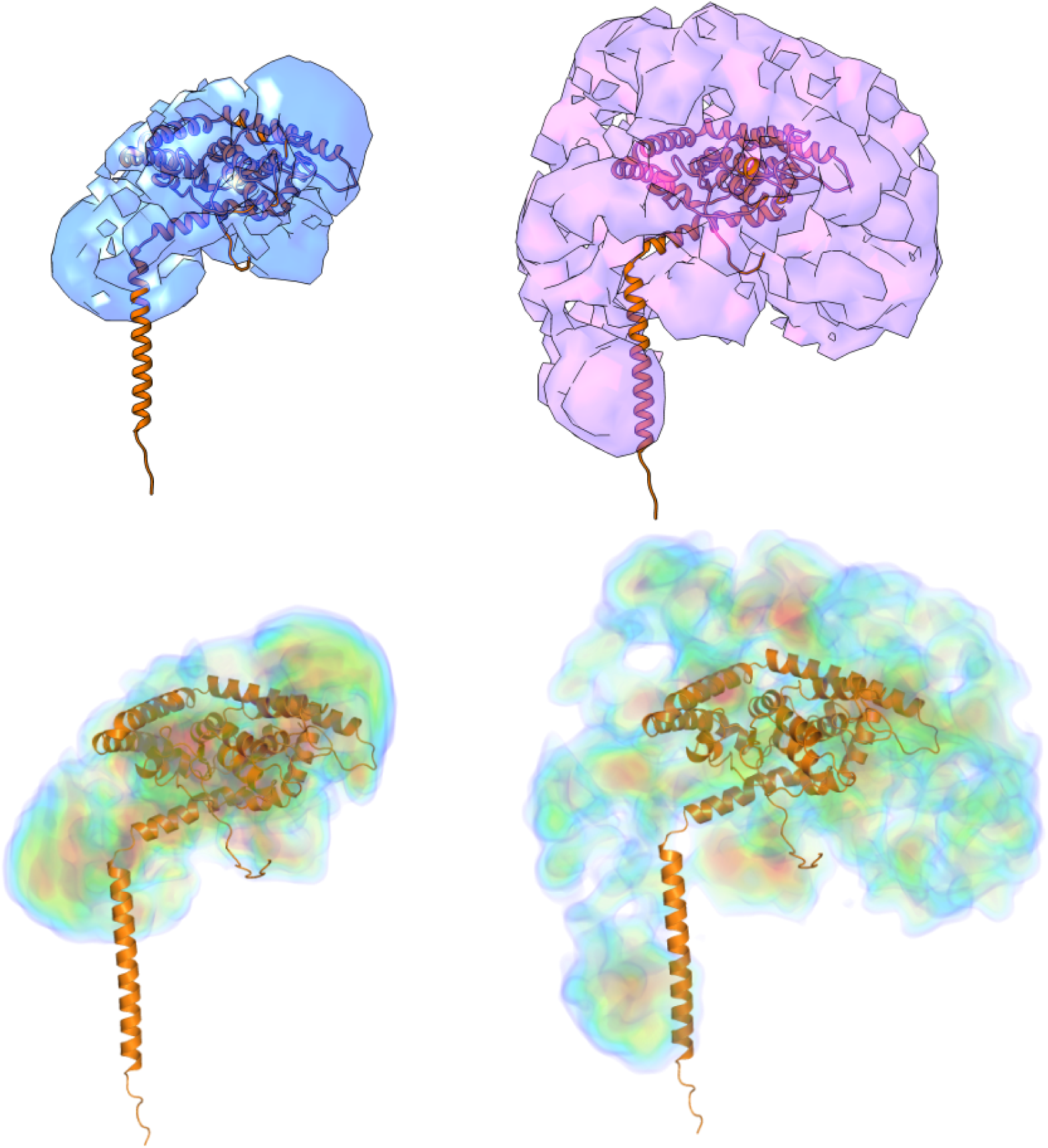
DENSS electron density models. Maps are shown for full-length GDAP1 without (left, blue) and with (right, purple) bound micelle. Superimposed (orange cartoon) is the full-length human GDAP1 AlphaFold2 model. The individual map reconstructions were iterated 20 times followed by averaging, alignment, and final map refinement. The rendering of the map contour (top) is set to the volume corresponding to the Porod volume (Å^2^). The density heat map (lower panel) is scaled with a relative contour lever based on the voxel volume.

**Figure S5.**
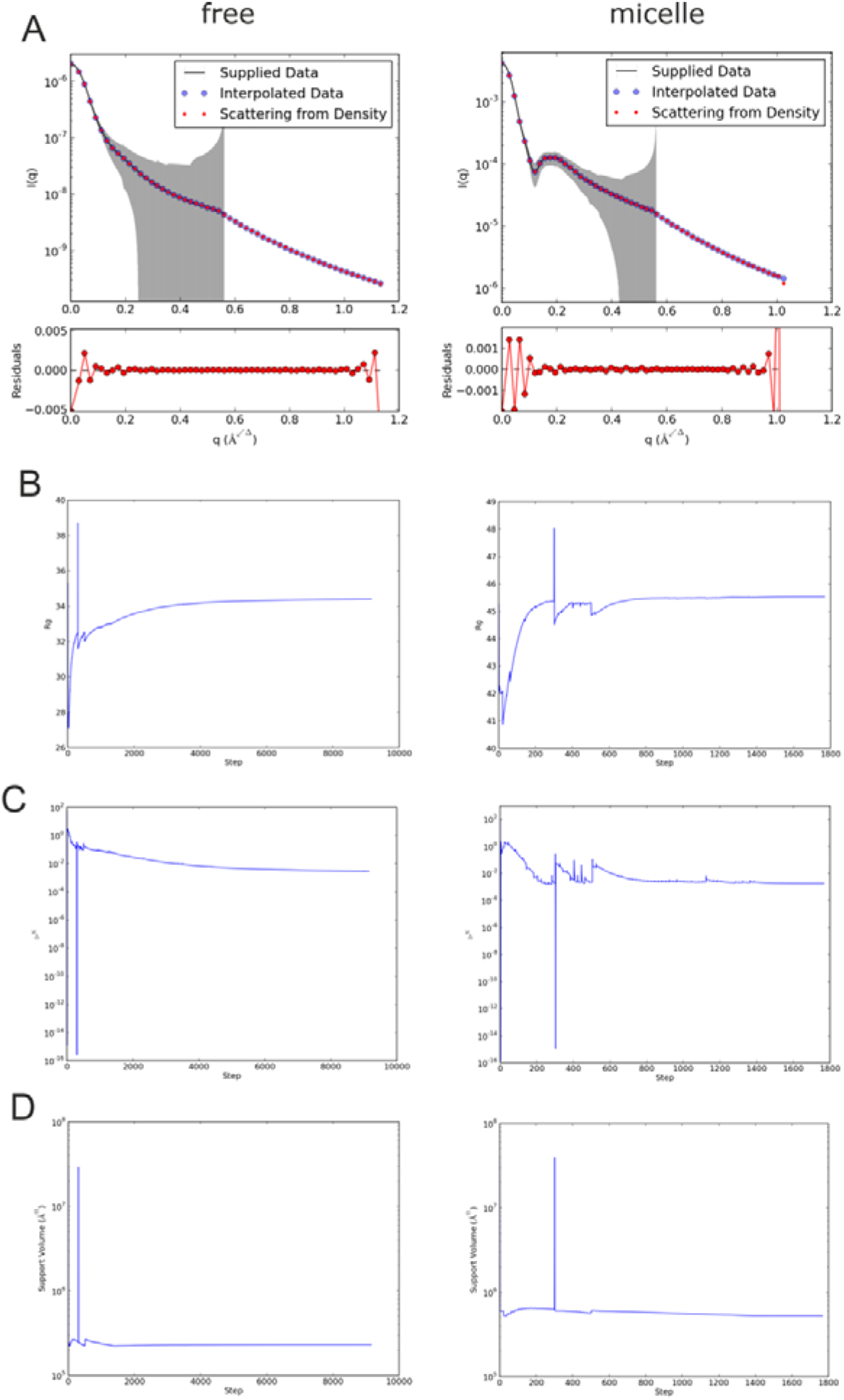
Reconstructed electron density map quality assessments of the final averaged and refined electron density maps from DENSS. A. Experimental SAXS scattering fit against the reconstructed scattering density. B-D. Development of R_g_ (B), chi^2^ (C), and support volume (D) over the course of the reconstruction iterative steps. In all panels, GDAP1 without micelle is on the left and the protein with bound micelle on the right.

**Figure S6.**
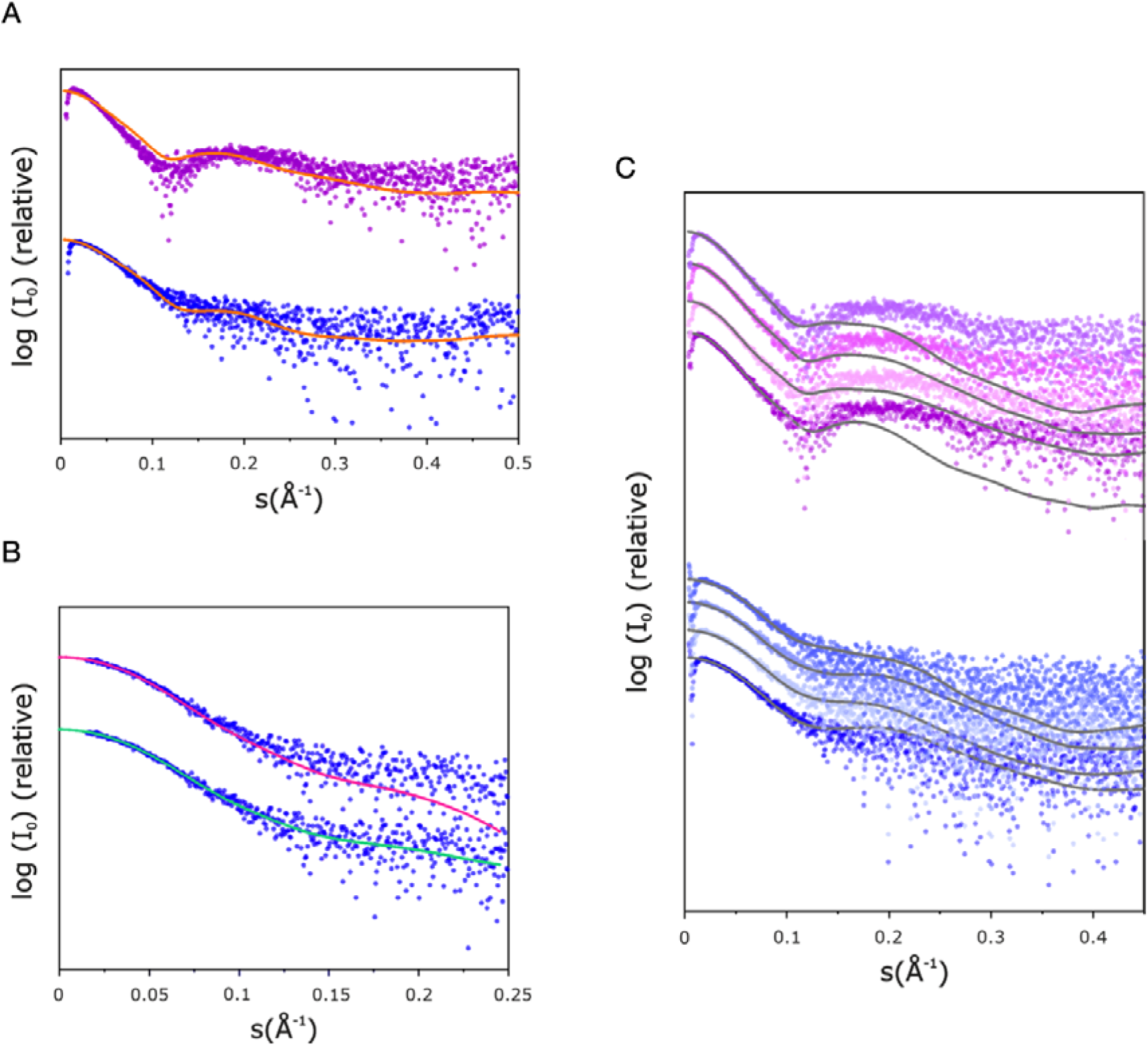
Fits of the SAXS models to experimental data. The SAXS data are coloured purple (first peak, with micelle) and blue (second peak, without micelle). A. Fit of the human GDAP1 AlphaFold2 model. B. Fit of the DAMMIN models. Note that in the presence of a micelle, DAMMIN fails to build a sensible model. C. Fits of the MONSA two-phase model to both datasets. MONSA was run 4 times, with different contrast values for the micelle phase.

## REFERENCES

1. Kann O, Kovacs R. Mitochondria and neuronal activity. Am J Physiol Cell Physiol. 2007;292(2):C641–57. Epub 20061108. doi: 10.1152/ajpcell.00222.2006. PubMed PMID: 17092996.

2. Rangaraju V, Lewis TL, Jr., Hirabayashi Y, Bergami M, Motori E, Cartoni R, et al. Pleiotropic Mitochondria: The Influence of Mitochondria on Neuronal Development and Disease. J Neurosci. 2019;39(42):8200–8. doi: 10.1523/JNEUROSCI.1157-19.2019. PubMed PMID: 31619488; PubMed Central PMCID: PMCPMC6794931.

3. Watts ME, Pocock R, Claudianos C. Brain Energy and Oxygen Metabolism: Emerging Role in Normal Function and Disease. Front Mol Neurosci. 2018;11:216. Epub 20180622. doi: 10.3389/fnmol.2018.00216. PubMed PMID: 29988368; PubMed Central PMCID: PMCPMC6023993.

4. Yin H, Zhu M. Free radical oxidation of cardiolipin: chemical mechanisms, detection and implication in apoptosis, mitochondrial dysfunction and human diseases. Free Radic Res. 2012;46(8):959–74. Epub 20120410. doi: 10.3109/10715762.2012.676642. PubMed PMID: 22468920.

5. Giacomello M, Pyakurel A, Glytsou C, Scorrano L. The cell biology of mitochondrial membrane dynamics. Nat Rev Mol Cell Biol. 2020;21(4):204–24. Epub 20200218. doi: 10.1038/s41580-020-0210-7. PubMed PMID: 32071438.

6. Niemann A, Wagner KM, Ruegg M, Suter U. GDAP1 mutations differ in their effects on mitochondrial dynamics and apoptosis depending on the mode of inheritance. Neurobiol Dis. 2009;36(3):509–20. Epub 20090925. doi: 10.1016/j.nbd.2009.09.011. PubMed PMID: 19782751.

7. Baxter RV, Ben Othmane K, Rochelle JM, Stajich JE, Hulette C, Dew-Knight S, et al. Ganglioside-induced differentiation-associated protein-1 is mutant in Charcot-Marie-Tooth disease type 4A/8q21. Nature genetics. 2002;30(1):21–2. doi: 10.1038/ng796.

8. Cuesta A, Pedrola L, Sevilla T, Garcia-Planells J, Chumillas MJ, Mayordomo F, et al. The gene encoding ganglioside-induced differentiation-associated protein 1 is mutated in axonal Charcot-Marie-Tooth type 4A disease. Nat Genet. 2002;30(1):22–5. Epub 20011217. doi: 10.1038/ng798. PubMed PMID: 11743580.

9. Marco A, Cuesta A, Pedrola L, Palau F, Marin I. Evolutionary and structural analyses of GDAP1, involved in Charcot-Marie-Tooth disease, characterize a novel class of glutathione transferase-related genes. Molecular biology and evolution. 2004;21(1):176–87. doi: 10.1093/molbev/msh013.

10. Rzepnikowska W, Kochanski A. A role for the GDAP1 gene in the molecular pathogenesis of CharcotMarieTooth disease. Acta neurobiologiae experimentalis. 2018;78(1):1–13.

11. Niemann A, Huber N, Wagner KM, Somandin C, Horn M, Lebrun-Julien F, et al. The Gdap1 knockout mouse mechanistically links redox control to Charcot-Marie-Tooth disease. Brain : a journal of neurology. 2014;137(Pt 3):668–82. doi: 10.1093/brain/awt371.

12. Niemann A, Ruegg M, La Padula V, Schenone A, Suter U. Ganglioside-induced differentiation associated protein 1 is a regulator of the mitochondrial network: new implications for Charcot-Marie-Tooth disease. The Journal of cell biology. 2005;170(7):1067–78. doi: 10.1083/jcb.200507087.

13. Pedrola L, Espert A, Wu X, Claramunt R, Shy ME, Palau F. GDAP1, the protein causing Charcot-Marie-Tooth disease type 4A, is expressed in neurons and is associated with mitochondria. Hum Mol Genet. 2005;14(8):1087–94. Epub 20050316. doi: 10.1093/hmg/ddi121. PubMed PMID: 15772096.

14. Barneo-Munoz M, Juarez P, Civera-Tregon A, Yndriago L, Pla-Martin D, Zenker J, et al. Lack of GDAP1 induces neuronal calcium and mitochondrial defects in a knockout mouse model of charcot-marie-tooth neuropathy. PLoS genetics. 2015;11(4):e1005115. doi: 10.1371/journal.pgen.1005115.

15. Cantarero L, Juárez-Escoto E, Civera-Tregón A, Rodríguez-Sanz M, Roldán M, Benítez R, et al. Mitochondria-lysosome membrane contacts are defective in GDAP1-related Charcot-Marie-Tooth disease. Human Molecular Genetics. 2021;29(22):3589–605. doi: 10.1093/hmg/ddaa243.

16. Miressi F, Benslimane N, Favreau F, Rassat M, Richard L, Bourthoumieu S, et al. GDAP1 Involvement in Mitochondrial Function and Oxidative Stress, Investigated in a Charcot-Marie-Tooth Model of hiPSCs-Derived Motor Neurons. Biomedicines. 2021;9(8). Epub 20210802. doi: 10.3390/biomedicines9080945. PubMed PMID: 34440148; PubMed Central PMCID: PMCPMC8393985.

17. Pijuan J, Cantarero L, Natera-de Benito D, Altimir A, Altisent-Huguet A, Diaz-Osorio Y, et al. Mitochondrial Dynamics and Mitochondria-Lysosome Contacts in Neurogenetic Diseases. Front Neurosci. 2022;16:784880. Epub 20220131. doi: 10.3389/fnins.2022.784880. PubMed PMID: 35177962; PubMed Central PMCID: PMCPMC8844575.

18. Wolf C, Lopez Del Amo V, Arndt S, Bueno D, Tenzer S, Hanschmann EM, et al. Redox Modifications of Proteins of the Mitochondrial Fusion and Fission Machinery. Cells. 2020;9(4). Epub 20200327. doi: 10.3390/cells9040815. PubMed PMID: 32230997; PubMed Central PMCID: PMCPMC7226787.

19. Wolf C, Pouya A, Bitar S, Pfeiffer A, Bueno D, Rojas-Charry L, et al. GDAP1 loss of function inhibits the mitochondrial pyruvate dehydrogenase complex by altering the actin cytoskeleton. Commun Biol. 2022;5(1):541. Epub 20220603. doi: 10.1038/s42003-022-03487-6. PubMed PMID: 35662277.

20. Sutinen A, Paffenholz D, Nguyen GTT, Ruskamo S, Torda AE, Kursula P. Conserved intramolecular networks in GDAP1 are closely connected to CMT-linked mutations and protein stability. PLoS One. 2023;18(4):e0284532. Epub 20230414. doi: 10.1371/journal.pone.0284532. PubMed PMID: 37058526; PubMed Central PMCID: PMCPMC10104300.

21. Googins MR, Woghiren-Afegbua AO, Calderon M, St. Croix CM, Kiselyov KI, VanDemark AP. Structural and functional divergence of GDAP1 from the glutathione S-transferase superfamily. The FASEB Journal. 2020;n/a(n/a). doi: 10.1096/fj.202000110R.

22. Nguyen GTT, Sutinen A, Raasakka A, Muruganandam G, Loris R, Kursula P. Structure of the Complete Dimeric Human GDAP1 Core Domain Provides Insights into Ligand Binding and Clustering of Disease Mutations. Frontiers in Molecular Biosciences. 2020;7:631232. doi: 10.3389/fmolb.2020.631232.

23. Sutinen A, Nguyen GTT, Raasakka A, Muruganandam G, Loris R, Ylikallio E, et al. Structural insights into Charcot-Marie-Tooth disease-linked mutations in human GDAP1. FEBS Open Bio. 2022. Epub 20220504. doi: 10.1002/2211-5463.13422. PubMed PMID: 35509130.

24. Moffitt W, Fitts DD, Kirkwood JG. Critique of the Theory of Optical Activity of Helical Polymers. Proc Natl Acad Sci U S A. 1957;43(8):723–30. doi: 10.1073/pnas.43.8.723. PubMed PMID: 16590076; PubMed Central PMCID: PMCPMC528528.

25. Burck J, Wadhwani P, Fanghanel S, Ulrich AS. Oriented Circular Dichroism: A Method to Characterize Membrane-Active Peptides in Oriented Lipid Bilayers. Acc Chem Res. 2016;49(2):184–92. Epub 20160112. doi: 10.1021/acs.accounts.5b00346. PubMed PMID: 26756718.

26. Krogh A, Larsson B, von Heijne G, Sonnhammer EL. Predicting transmembrane protein topology with a hidden Markov model: application to complete genomes. J Mol Biol. 2001;305(3):567–80. doi: 10.1006/jmbi.2000.4315. PubMed PMID: 11152613.

27. Jumper J, Evans R, Pritzel A, Green T, Figurnov M, Ronneberger O, et al. Highly accurate protein structure prediction with AlphaFold. Nature. 2021;596(7873):583-9. Epub 20210715. doi: 10.1038/s41586-021-03819-2. PubMed PMID: 34265844; PubMed Central PMCID: PMCPMC8371605.

28. Beaugrand M, Arnold AA, Henin J, Warschawski DE, Williamson PT, Marcotte I. Lipid concentration and molar ratio boundaries for the use of isotropic bicelles. Langmuir. 2014;30(21):6162–70. Epub 20140519. doi: 10.1021/la5004353. PubMed PMID: 24797658; PubMed Central PMCID: PMCPMC4072726.

29. Chen FY, Lee MT, Huang HW. Sigmoidal concentration dependence of antimicrobial peptide activities: a case study on alamethicin. Biophys J. 2002;82(2):908–14. doi: 10.1016/S0006-3495(02)75452-0. PubMed PMID: 11806932; PubMed Central PMCID: PMCPMC1301899.

30. Miles AJ, Wallace BA. CDtoolX, a downloadable software package for processing and analyses of circular dichroism spectroscopic data. Protein science : a publication of the Protein Society. 2018;27(9):1717–22. doi: 10.1002/pro.3474.

31. Thureau A, Roblin P, Pérez J. BioSAXS on the SWING beamline at Synchrotron SOLEIL. J Appl Crystallogr. 2021;54:1698–710.

32. Manalastas-Cantos K, Konarev PV, Hajizadeh NR, Kikhney AG, Petoukhov MV, Molodenskiy DS, et al. ATSAS 3.0: expanded functionality and new tools for small-angle scattering data analysis. J Appl Crystallogr. 2021;54(Pt 1):343–55. Epub 20210201. doi: 10.1107/S1600576720013412. PubMed PMID: 33833657; PubMed Central PMCID: PMCPMC7941305.

33. Maupetit J, Derreumaux P, Tuffery P. PEP-FOLD: an online resource for de novo peptide structure prediction. Nucleic Acids Res. 2009;37(Web Server issue):W498-503. Epub 20090511. doi: 10.1093/nar/gkp323. PubMed PMID: 19433514; PubMed Central PMCID: PMCPMC2703897.

34. Lomize AL, Todd SC, Pogozheva ID. Spatial arrangement of proteins in planar and curved membranes by PPM 3.0. Protein Sci. 2022;31(1):209–20. Epub 20211108. doi: 10.1002/pro.4219. PubMed PMID: 34716622; PubMed Central PMCID: PMCPMC8740824.

35. Jo S, Cheng X, Lee J, Kim S, Park SJ, Patel DS, et al. CHARMM-GUI 10 years for biomolecular modeling and simulation. J Comput Chem. 2017;38(15):1114–24. Epub 20161114. doi: 10.1002/jcc.24660. PubMed PMID: 27862047; PubMed Central PMCID: PMCPMC5403596.

36. Grant TD. Ab initio electron density determination directly from solution scattering data. Nat Methods. 2018;15(3):191–3. Epub 20180129. doi: 10.1038/nmeth.4581. PubMed PMID: 29377013.

37. Raasakka A, Ruskamo S, Kowal J, Han H, Baumann A, Myllykoski M, et al. Molecular structure and function of myelin protein P0 in membrane stacking. Sci Rep. 2019;9(1):642. Epub 20190124. doi: 10.1038/s41598-018-37009-4. PubMed PMID: 30679613; PubMed Central PMCID: PMCPMC6345808.

38. Opalinski L, Becker T, Pfanner N. Clearing tail-anchored proteins from mitochondria. Proc Natl Acad Sci U S A. 2014;111(22):7888–9. Epub 20140522. doi: 10.1073/pnas.1406864111. PubMed PMID: 24853504; PubMed Central PMCID: PMCPMC4050541.

39. Rojo M, Legros F, Chateau D, Lombes A. Membrane topology and mitochondrial targeting of mitofusins, ubiquitous mammalian homologs of the transmembrane GTPase Fzo. J Cell Sci. 2002;115(Pt 8):1663–74. doi: 10.1242/jcs.115.8.1663. PubMed PMID: 11950885.

40. Wagner KM, Ruegg M, Niemann A, Suter U. Targeting and function of the mitochondrial fission factor GDAP1 are dependent on its tail-anchor. PloS one. 2009;4(4):e5160. doi: 10.1371/journal.pone.0005160.

41. Azzedine H, Ruberg M, Ente D, Gilardeau C, Perie S, Wechsler B, et al. Variability of disease progression in a family with autosomal recessive CMT associated with a S194X and new R310Q mutation in the GDAP1 gene. Neuromuscul Disord. 2003;13(4):341–6. PubMed PMID: 12868504.

42. Rodriguez-Hernandez A, Mayo M, Jauregui L, Patel P. Autosomal dominant GDAP1 mutation with severe phenotype and respiratory involvement: A case report. Front Neurol. 2022;13:905725. Epub 20221024. doi: 10.3389/fneur.2022.905725. PubMed PMID: 36353131; PubMed Central PMCID: PMCPMC9637907.

43. Yoshimura A, Yuan JH, Hashiguchi A, Hiramatsu Y, Ando M, Higuchi Y, et al. Clinical and mutational spectrum of Japanese patients with Charcot-Marie-Tooth disease caused by GDAP1 variants. Clin Genet. 2017;92(3):274–80. Epub 20170419. doi: 10.1111/cge.13002. PubMed PMID: 28244113.

44. Alfares A, Alfadhel M, Wani T, Alsahli S, Alluhaydan I, Al Mutairi F, et al. A multicenter clinical exome study in unselected cohorts from a consanguineous population of Saudi Arabia demonstrated a high diagnostic yield. Mol Genet Metab. 2017;121(2):91–5. Epub 20170407. doi: 10.1016/j.ymgme.2017.04.002. PubMed PMID: 28454995.

45. van Paassen BW, Bronk M, Verhamme C, van Ruissen F, Baas F, van Spaendonck-Zwarts KY, et al. Pseudodominant inheritance pattern in a family with CMT2 caused by GDAP1 mutations. J Peripher Nerv Syst. 2017;22(4):464–7. Epub 20170911. doi: 10.1111/jns.12236. PubMed PMID: 28837237.

46. Nykamp K, Anderson M, Powers M, Garcia J, Herrera B, Ho YY, et al. Sherloc: a comprehensive refinement of the ACMG-AMP variant classification criteria. Genet Med. 2017;19(10):1105–17. Epub 20170511. doi: 10.1038/gim.2017.37. PubMed PMID: 28492532; PubMed Central PMCID: PMCPMC5632818.

47. Sivera R, Sevilla T, Vilchez JJ, Martinez-Rubio D, Chumillas MJ, Vazquez JF, et al. Charcot-Marie-Tooth disease: genetic and clinical spectrum in a Spanish clinical series. Neurology. 2013;81(18):1617–25. Epub 20130927. doi: 10.1212/WNL.0b013e3182a9f56a. PubMed PMID: 24078732; PubMed Central PMCID: PMCPMC3806911.

48. Schenkel LC, Bakovic M. Formation and regulation of mitochondrial membranes. Int J Cell Biol. 2014;2014:709828. Epub 20140122. doi: 10.1155/2014/709828. PubMed PMID: 24578708; PubMed Central PMCID: PMCPMC3918842.

